# Receptor-like kinases FER and SRK mediate *Lotus japonicus* freezing tolerance and climate adaptation

**DOI:** 10.1101/2022.04.27.489728

**Authors:** Yusdar Mustamin, Turgut Yigit Akyol, Max Gordon, Andi Madihah Manggabarani, Yoshiko Isomura, Yasuko Kawamura, Masaru Bamba, Cranos Williams, Stig Uggerhøj Andersen, Shusei Sato

**Affiliations:** Graduate School of Life Sciences, Tohoku University, Katahira 2-1-1, Aoba-ku, Sendai, 980-8577, Japan; Department of Molecular Biology and Genetics, Aarhus University, DK-8000 Aarhus, Denmark; Department of Electrical and Computer Engineering, North Carolina State University, 890 Oval Drive, 3114 Engineering Building II, Raleigh, NC 27606, USA

**Keywords:** Climate adaptation, cold acclimation, freezing tolerance, legume, perennial persistence, RNA-seq

## Abstract

Many plant species have succeeded in colonizing a wide range of diverse climates through local adaptation, but the underlying molecular genetics remain obscure. We previously found that winter survival was a direct target of selection during colonization of Japan by the perennial legume *Lotus japonicus* and identified associated candidate genes. Here, we show that two of these, the *FERONIA-receptor like kinase* (*LjFER*) and a novel S-receptor-like kinase (*LjSRK*) are required for non-acclimated freezing tolerance and show haplotype-dependent cold-responsive expression. Our work demonstrates that recruiting a conserved growth regulator, *FER*, and a novel receptor-like kinase, *SRK*, into the set of cold-responsive genes has contributed to freezing tolerance and local climate adaptation in *L. japonicus*, offering new functional genetic insight into perennial herb evolution.

## INTRODUCTION

Global cooling events resulted in establishment of temperate regions with seasonal temperature fluctuations about 47 to 26 million years ago. A large number of plant families subsequently adapted to and colonized these new environments. This process involved a number of independent events across many lineages, suggesting that there were limited requirements for pre-adaptation and that it was relatively unproblematic to adapt to cold winters by recruiting genetic components from existing signaling pathways such as drought tolerance (Preston and Sandve, 2013). Some of the most important crops, including winter wheat, are winter annuals that are sown and germinate in autumn and flower and set seed in spring and summer the following year. Together, the evolutionary history and importance for crop productivity have spurred a strong research interest in freezing tolerance and adaptation to temperate climates.

The molecular genetics of freezing tolerance has been most extensively studied in *Arabidopsis thaliana* (Arabidopsis), which includes both summer and winter annuals (Koornneef et al., 2010). Research efforts were mainly focused on cold acclimation, a process where low, non-freezing temperatures pre-condition plants to better cope with exposure to frost (Thomashow, 1999). A large transcriptional response was described as a component of cold acclimation, and the *CBF* genes were identified as important players in the control of a set of cold regulated genes (Gilmour and Thomashow, 1991; Gilmour et al., 1998). Additionally, the ICE1 transcription factor was identified as a regulator of the *CBF* genes (Kurbidaeva et al., 2015).

Also in Arabidopsis, a considerable effort was made to determine the genetic components contributing to cold adaptation through natural variation and selection across Arabidopsis accessions. Mapping using biparental populations identified several quantitative trait loci (QTLs), some of which overlapped with the *CBF* genes (Alonso-Blanco et al., 2005; Zhen and Ungerer, 2008a; Kang et al., 2013; Oakley et al., 2014; Gehan et al., 2015). *CBF2* was identified as a causal gene underlying a specific QTL, where the southern parental accession carried a loss-of-function allele (Alonso-Blanco et al., 2005). Other studies did not find direct evidence of CBF importance for natural variation in freezing tolerance (Le et al., 2008; Suther et al., 2012; Meissner et al., 2013), suggesting that both CBF-dependent and - independent adaptation has occurred. Arabidopsis natural variation for freezing tolerance was also explored in a large genome-wide association (GWA) study using nearly 500 global accessions. A number of candidate genes were identified, including cold-regulated, but not *CBF* genes (Horton et al., 2016). In other species, QTL overlapping *CBF* genes have also been identified, but a causal role has not been established (Knox et al., 2010; Tayeh et al., 2013; Würschum et al., 2017).

Freezing tolerance is classified as cold-acclimated, following exposure to low, non-freezing temperatures, or non-acclimated. Non-acclimated freezing tolerance has been less extensively studied than its cold-acclimated counterpart, but there is evidence for correlation between the two traits, suggesting an overlap in genetic components (Zhen and Ungerer, 2008b). This is supported by the identification of the *M. truncatula* receptor-like kinase *MtCTLK* as important for both cold-acclimated and non-acclimated freezing tolerance and full *CBF* gene induction (Geng et al., 2021). The relative adaptive importance of cold-acclimated and non-acclimated freezing tolerance for adaptation remains an open question, and it is likewise unclear what trajectories adaptation to colder climates has followed in perennial herbs.

*Lotus japonicus* (Lotus) is a perennial herbaceous legume that grows across a wide range of Japanese geographical locations and climate conditions from the subtropical south to the temperate north, and from coastal areas to alpine environments. Unlike winter annuals, such as those found in Arabidopsis and wheat, northern Lotus accessions have no vernalization requirement and shed their leaves in autumn followed by regrowth from crown buds the following spring. In contrast, southern Lotus accessions stay green throughout the year, allowing more rapid accumulation of biomass in the absence of low winter temperatures. The ubiquitous presence across climatic gradients combined with a well-described colonization history and a clear population structure with southern (pop1), central (pop2) and northern (pop3) populations, makes Lotus well suited for studying perennial herb climate adaptation (Shah et al., 2020). Using a combination of genome-wide association (GWA) scans and population differentiation analysis, we previously identified in-field winter survival as the trait with the strongest selective signature during *Lotus japonicus* colonization of Japan and discovered associated candidate genes (Shah et al., 2020). Here, we carry out a detailed investigation of two of these, the receptor-like kinases *LjFER* and *LjSRK*, to demonstrate that they are required for non-acclimated freezing tolerance and display haplotype-dependent cold responsiveness.

## RESULTS

### Transcriptional population differentiation identifies overwintering candidate genes

Using the Lotus MG20 v.3.0 reference genome, we had previously detected several overwintering-associated genomic regions based on data from field experiments carried out in 2014 (Shah et al., 2020). Here, we repeated the GWA analysis using the new Gifu v.1.2 reference assembly (Kamal et al., 2020) (Supplemental Table S1) and identified signals consistent with our previous analysis. Some of the identified regions were relatively broad and contained many candidate genes (Figure 1A; Supplemental Table S2), making it challenging to pinpoint specific genes for in-depth investigation.

**Figure 1.**
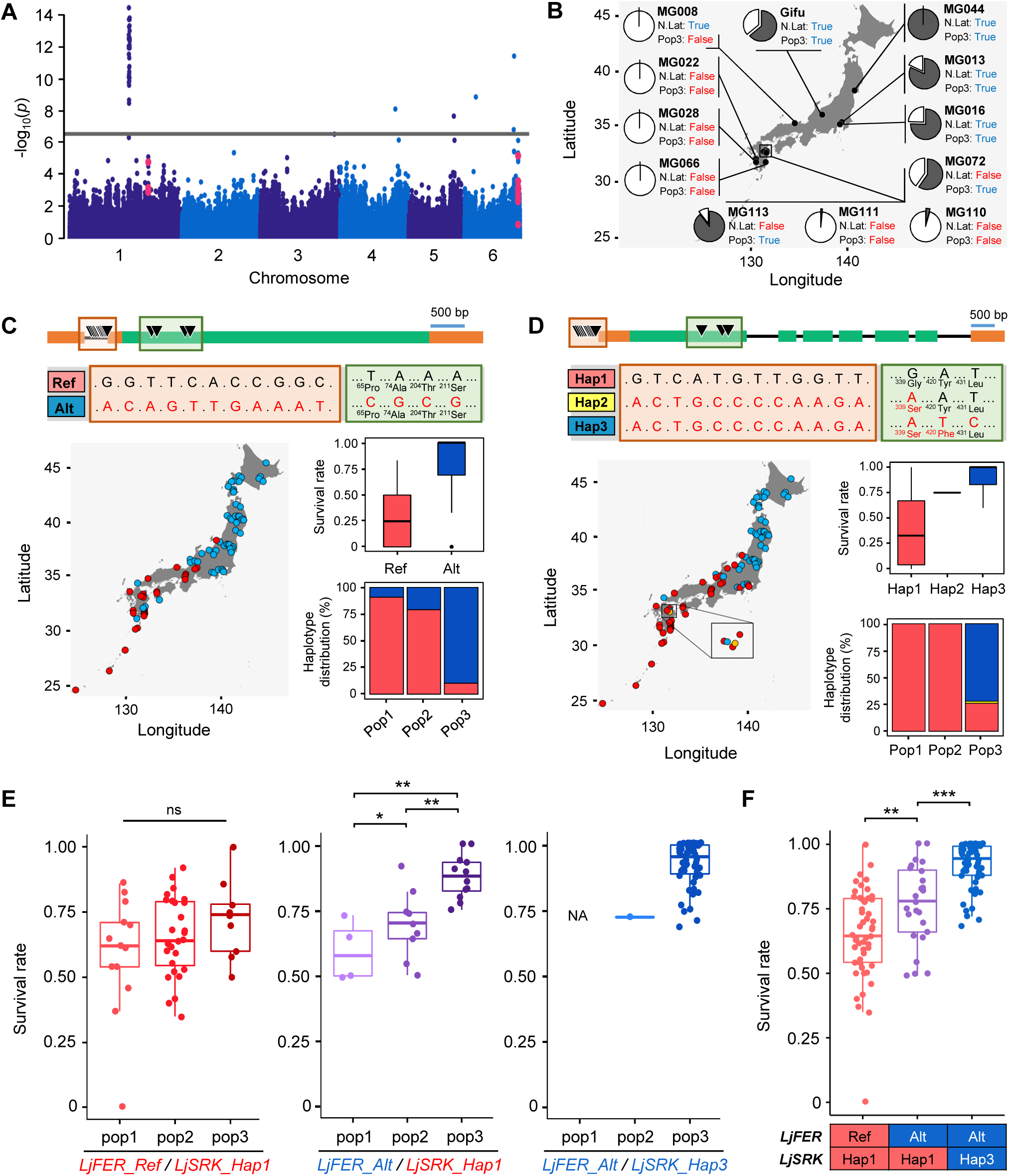
Winter-survival candidate genes. A) Manhattan plot of a GWA scan for winter survival in 2014 at the Tohoku field site. *LjFER* and *LjSRK* SNPs are highlighted (red dots). B) Origins of the twelve accessions selected for seasonal transcriptional profiling. Pie charts indicate their pop3 membership (dark gray slice). Their Northern latitude (N. lat) and Pop3 membership (Pop3) binary classification in the logistic regression analysis is indicated. C) *LjFER* haplotype distribution. D) *LjSRK* haplotype distribution. Red letters indicate differences with respect to the Gifu haplotype (Ref or Hap1). E-F) Winter survival in 2014 for accessions split by the *LjFER* and *LjSRK* haplotypes or haplotype combinations indicated (One-way ANOVA with posthoc Tukey’s test: ns = no significant (*P* > 0.05), * = *P* < 0.05, ** = *P* < 0.01, *** = *P* < 0.001).

In our earlier study, we had observed pronounced genetic differentiation at specific loci between the northern *L. japonicus* population, pop3, versus the southern (pop1) and central (pop2) populations at the same SNPs that showed association with winter survival (Shah et al., 2020). We reasoned that transcriptional profiles could similarly be queried for differentiation signals in order to provide us with single-gene resolution for candidate gene selection. To generate data for this transcriptome analysis, we selected a subset of Japanese *L. japonicus* accessions designed to facilitate disentangling geographic origin and population structure (Figure 1B). This subset included southern accessions, which showed an unusually high level of pop3 membership, MG072 and MG113, as well as a northern accession that did not belong to pop3, MG008 (Figure 1B).

We planted the accessions in a field located near Tohoku University and carried out transcriptional profiling of leaf samples collected in August, October, December and February 2018/2019 (Supplemental Table S3). We divided the accessions into two groups both with respect to geographical origin and pop3 membership (Figure 1B) and then used logistic regression to evaluate how well the accessions could be separated with respect to these two traits based on the seasonal expression profile of a given gene. There were 3164 genes where a maximum of one accession was misclassified for pop3 membership. Out of those, 1901 genes showed better classification of pop3 membership than geographical origin (Supplemental Table S4). Intersecting our GWA results with the 1901 candidate genes from the logistic regression analysis left 59 SNPs of which nine out of the top ten were associated with two receptor-like kinases (Supplemental Table S5). The first was the G-type lectin S-receptor-like serine/threonine-protein kinase, *LotjaGi6g1v0343100*, which we name *LjSRK*. The second was the multifunctional receptor-like kinase *FERONIA (LjFER), LotjaGi1g1v0458500*. Based on the combined GWA and transcriptome analysis, we chose to focus on these two candidate genes.

### *LjFER* and *LjSRK* haplotypes are associated with variation in winter survival

*LjFER* showed 11 SNPs in an intron within the 5’UTR and four silent SNPs in the coding region (Figure 1C). There were two distinct *FER* haplotypes (Figure 1C). Although both haplotypes were present in all Lotus populations, *LjFER_Ref* was mainly found in pop1 (89.8%) and pop2 (77.3%), which generally showed low winter survival, while *LjFER_Alt* was found in 88.6% of the winter-hardy pop3 accessions. *LjSRK* showed a total of 15 SNPs, 12 in the promoter region and three in the coding region, and comprised three haplotypes. *LjSRK*_*Hap1* is the reference (Gifu) haplotype; *LjSRK*_*Hap2* has a single amino acid substitution with respect to the reference; and *LjSRK*_*Hap3* shows one additional amino acid change compared to *LjSRK_Hap2* (Figure 1D). There were no differences between *LjSRK*_*Hap2* and *LjSRK*_*Hap3* with respect to the promoter SNPs (Figure 1D).

To investigate the winter survival characteristics of the *LjFER* and *LjSRK* haplotypes in more detail, we re-examined data from the Tohoku field site in 2014 (Shah et al., 2020). Splitting the accessions by population and *LjFER* and *LjSRK* haplotypes, we observed no significant differences between populations for accessions with the *LjFER_Ref/LjSRK_Hap1* haplotype (Figure 1E). In contrast, *LjFER_Alt/LjSRK_Hap1* accessions showed significant survival differences between subpopulations (Figure 1E). The *LjFER_Alt/LjSRK_Hap3* combination was nearly exclusive to pop3 and the accessions with this haplotype showed the highest median winter survival (Figure 1E). The *LjFER_Ref/LjSRK_Hap3* haplotype was not found in our population sample. Across all populations, accessions carrying the *LjFER_Alt* allele showed increased survival (One-way ANOVA with posthoc Tukey’s test, *P <* 0.01) and there was a clear additive effect with *LjSRK_Hap3* (One-way ANOVA with posthoc Tukey’s test, *P <* 0.001) (Figure 1F).

Our earlier work indicated that winter survival in field experiments depended greatly on the conditions of each specific year (Shah et al., 2020). In addition, experimental replication was limited to one winter climate per site per year. To establish a more flexible experimental system that would allow assaying multiple different cold challenges per year in the absence of cold acclimation, we grew Lotus accessions in the glasshouse and transferred them to unprotected outdoor conditions for a four week period before returning them to the glasshouse to recover. Using this approach, we exposed plants to shocks of different severity, depending on the timing of the outdoor transfer. We examined 23 Lotus accessions selected based on the haplotype combinations of *LjFER* and *LjSRK* and geographical origin (Figure 2A). The first round of experiments was carried out in winter 2019/20, when the lowest temperature -4.7°C was reached in mid-February (W7) (Figure 2B). In winter 2020/21, the minimum temperature of -7.6°C occurred in early January (W3) (Figure 2C). The survival rates tracked the temperature curves during the test period in both experimental years (Figure 2, B and C; Supplemental Table S6), indicating that the outdoor temperature during the challenge influenced the survival rates for all accessions. Again, we observed pronounced differences in survival between the different *LjFER/LjSRK* haplotypes and we found the highest survival rates in the *LjFER_Alt/LjSRK_Hap3* accessions (Figure 2, B and C).

**Figure 2.**
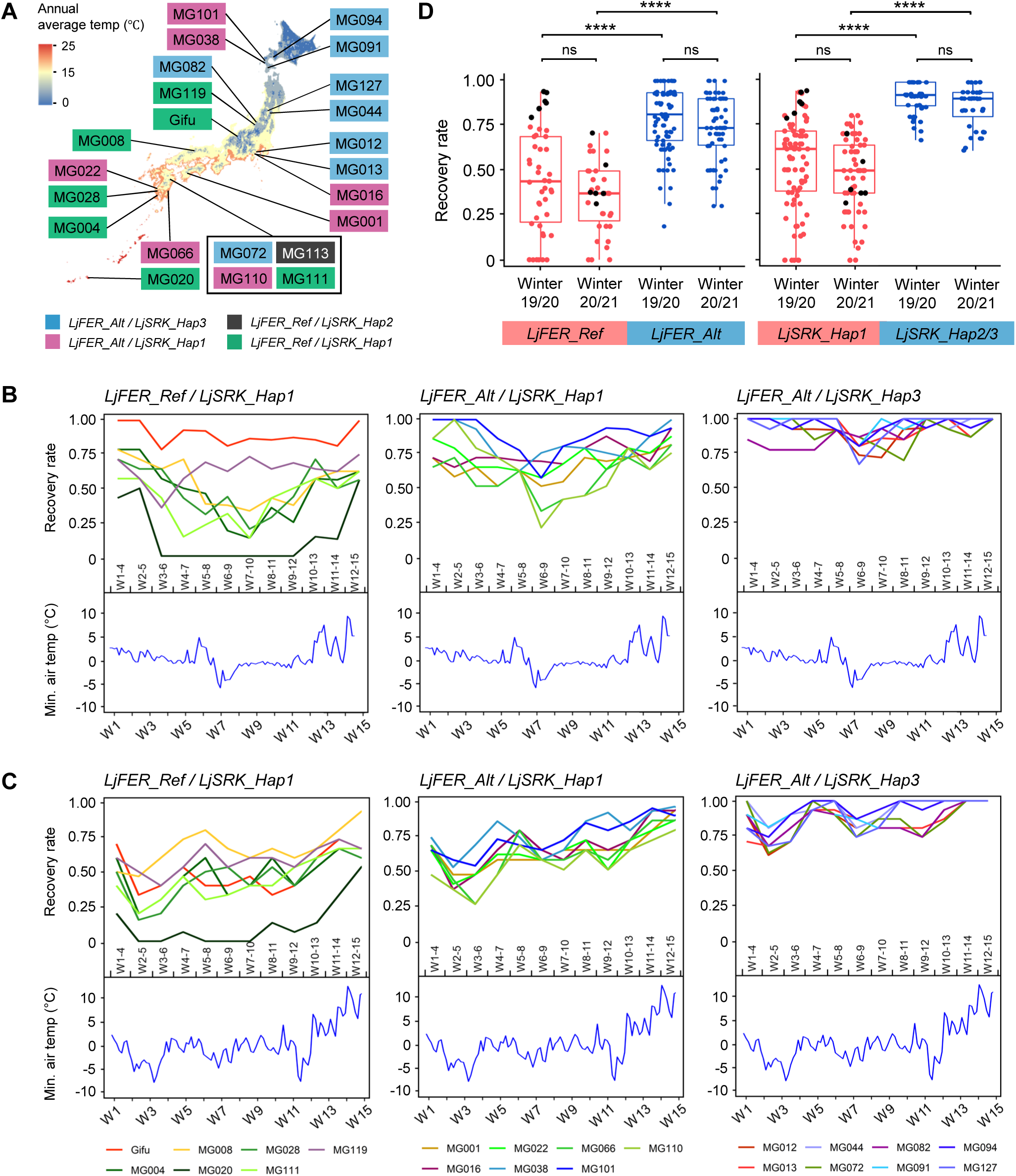
Glasshouse experiment with outdoor winter challenge. A) Origins and *LjFER*/*LjSRK* haplotypes of the 23 accessions selected for the glasshouse experiments. B-C) Accession recovery rate and outdoor air temperature winter 2019/20 (B) and 2020/21 (C). The period of the outdoor challenge is indicated at the bottom of the upper panel. E.g. W1-4 indicates that the plants were exposed to outdoor winter conditions in weeks one through four in 2021. Survival rates were determined after two weeks of recovery inside the glasshouse following the outdoor exposure. The x-axis on the lower panels indicates the week number in which the temperature was measured. (n=13-16). D) Boxplots showing recovery rate grouped by *LjSRK* and *LjFER* haplotypes. Each dot represents the recovery rate when the temperature minimum occurred (W7 to W12 in winter 2019/20 and W1 to W6 in winter 2020/21) (Paired student’s t-test: ns = no significant, **** = *P* < 0.0001). Data points from Gifu are highlighted in black.

With respect to the *LjFER/LjSRK* haplotype effects, we also observed a marked difference between years. In winter 2019/20, when the coldest period occurred relatively late, the *LjFER_Ref* and *LjSRK_Hap1* accessions showed some overlap in survival rate with the *LjFER_Alt* and *LjSRK_Hap2/3* accessions, some reaching survival rates of more than 90% (Figure 2, B-D). This overlap was reduced in winter 2020/21, where a stronger differentiation was observed between the accessions carrying different *LjFER/LjSRK* haplotypes (Figure 2, B-D). The trend was especially clear for Gifu, which originates from central Japan (Figure 2). It shows mixed pop2/3 membership and a *LjFER_Ref/LjSRK_Hap1* haplotype. In winter 2019/20, it showed the highest survival among the *LjFER_Ref/LjSRK_Hap1* accessions during the coldest period, whereas its survival was strongly reduced in 2020/2021 (Figure 2, B-D). This is consistent with the *LjFER_Alt/LjSRK_Hap3* haplotype being more critical for freezing tolerance under more challenging conditions.

### *LjFER* and *LjSRK* are required for freezing tolerance

The non-transgenic *LORE1* mutant resource was generated in the Gifu genetic background (Małolepszy et al., 2016). We therefore used *LORE1* mutants to test whether *LjFER* and *LjSRK* play a role in freezing tolerance, even in an accession that does not carry the *LjFER_Alt* and *LjSRK_Hap3* haplotypes. We identified a single heterozygous insertion line for *LjFER* and three homozygous insertion lines for *LjSRK* (Małolepszy et al., 2016; Mun et al., 2016) (Figure 3A). Since homozygous insertion lines of *fer-1* could not be obtained, we checked the expression level of *LjFER* in each heterozygous insertion line to examine the possibility that the heterozygous insertions might cause expression instability. We found that the gene expression level of *LjFER* was significantly different between the tested heterozygous *fer-1* individuals and was greatly reduced compared to the Gifu wild type (Supplemental Figure S1A). Therefore, we used heterozygous *fer-1* lines for further functional analyses. We found extremely reduced *LjSRK* expression levels in all three *LORE1* insertion lines (Supplemental Figure S1B).

**Figure 3.**
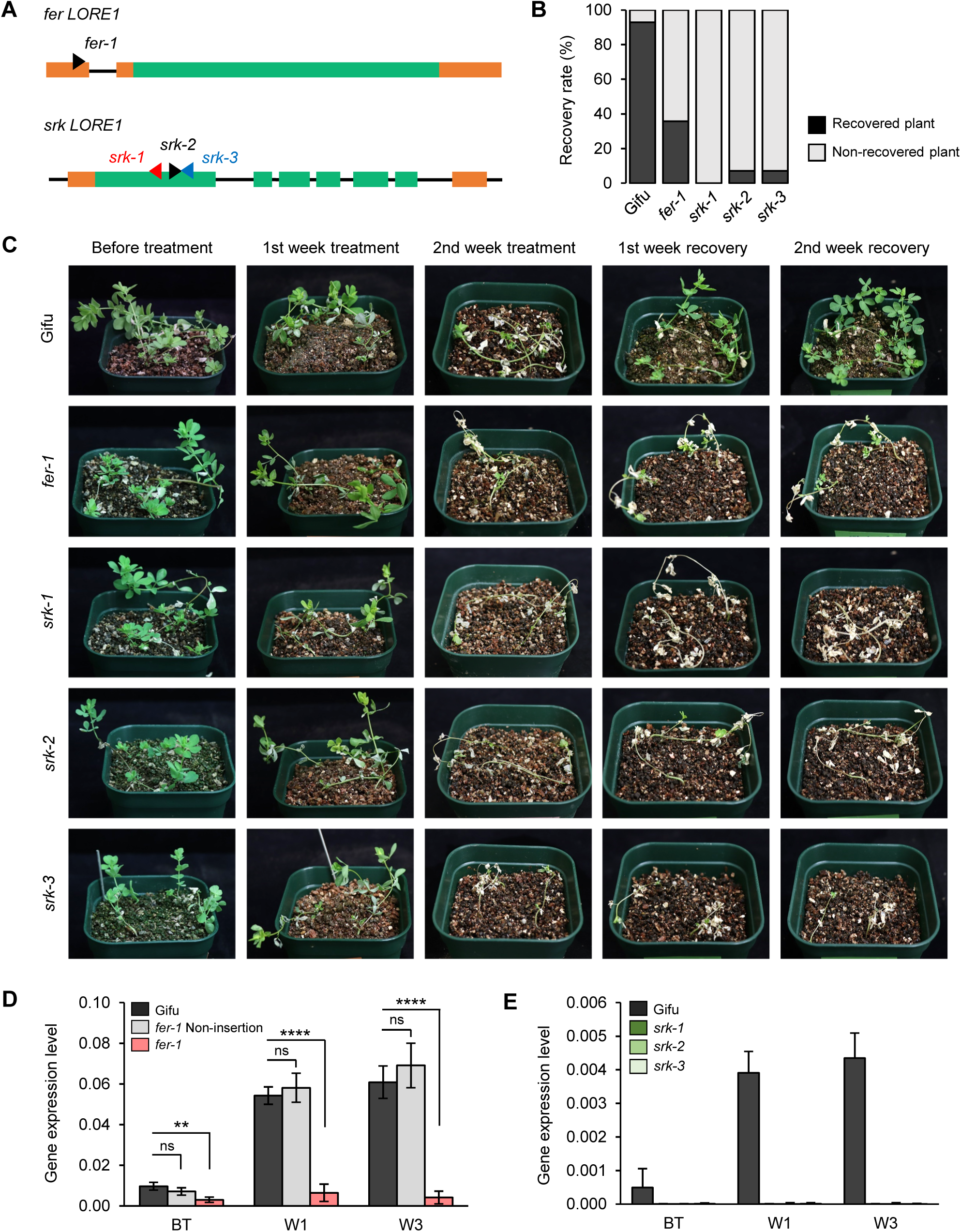
*fer* and *srk LORE1* mutants. A) *LORE1* insertion sites in *LjFER* (*fer-1*) and *LjSRK* (*srk-1,-2,-3*). B) Final recovery rates and, C) the phenotypes of the indicated genotypes in the 2019/20 glasshouse with outdoor exposure experiment (n = 14 each). D-E) *LjFER* (D) and *LjSRK* (E) induction by cold treatment in week-1 (W1) and week-3 (W3) of winter 2020/21 experiment (exp-1). The gene expression bars indicate the mean value ± SD of six biological replicates combined with three technical replicates (Paired student’s t-test: ns = no significant, ** = *P* < 0.01, **** = *P* < 0.0001). Additional experimental replicates are shown in Supplemental Figure S6.

We then grew the non-transgenic *LORE1* mutants at the Tohoku field site during summer and examined the overwintering rate in spring the following year. We found 15% to 30% reduction of winter survival in the *srk* mutants, whereas the *fer-1* mutants did not display reduced survival compared to the Gifu wild type (Supplemental Figure S1C). We then challenged the mutants using the outdoor exposure method described above (see the Materials and Methods section). The minimum air temperature during the four weeks of outdoor exposure reached -1.2°C and -7.6°C for the winter 2019/20 and winter 2020/21 experiments, respectively (Supplemental Figure S2, A and B). During winter 2019/20 (Figure 3C), all *LORE1* insertion lines and the Gifu wild type stayed green, albeit with some leaf shedding in the first week of treatment followed by severe damage with etiolated leaves in the fourth week of treatment. After one week of recovery, green leaves appeared in Gifu and in some *fer-1* individuals. These also exhibited new crown buds during their final recovery. We found extremely low recovery rates in all *LjSRK_LORE1* insertion lines compared to Gifu (Figure 3B). Since the phenotypic effects of the *fer-1* insertion were not as clear as for the *srk* insertions, we compared the expression level of *LjFER* in the first and third weeks of outdoor exposure with the survival phenotype. Individuals with poor recovery showed significantly lower *LjFER* expression in both weeks 1 and 3 (Supplemental Figure S3A), indicating that strong reductions in *LjFER* expression were detrimental to recovery from cold exposure. We repeated the *LORE1* mutant experiments in winter 2020/21 with consistent results (Supplemental Figures S4-S7). Furthermore, we carried out additional transcriptional profiling in 2020/21, which showed induction of *LjFER* and *LjSRK* during cold treatment in the Gifu wild type and absence of induction in the *fer-1* and *srk* mutants (Figure 3, D and E; Supplemental Figure S6, A and B).

### *LjFER* and *LjSRK* show haplotype-specific temperature-dependent expression

Gifu carries the *LjFER_Ref/LjSRK_Hap1* haplotype and the *LORE1* mutant experiments demonstrated that loss of expression of either of the two genes, despite their non-winter hardy haplotypes, was detrimental to freezing tolerance. In addition, *LjFER* lacked coding region SNPs and the logistic regression analysis of seasonal gene expression indicated that *LjFER* and *LjSRK* profiles could predict winter survival. To determine if the different *LjFER* and *LjSRK* alleles had distinct expression patterns, which could explain their effects on winter survival and freezing tolerance, we carried out qRT-PCR gene expression analysis using samples collected after the first week of cold exposure in the glasshouse experiments. For *LjFER_Ref* accessions, we observed a small increase in expression concomitant with a drop in outdoor air temperature (Figure 4, A and B). This trend was much more pronounced for *LjFER_Alt*, and the expression peak was clearly different between the two years, coinciding with the first major drop in temperature to below freezing (Figure 4, A and B). For both years, there were significant differences in expression between the *LjFER* haplotypes at the expression peaks in weeks 8 and 3, respectively (Paired Student’s t-test, *P <* 0.0001). A similar temperature-dependent response was seen in both years for *LjSRK_Hap3* accessions, whereas there was no indication of temperature-dependent expression changes for *LjSRK_Hap1*. Expression differences between accessions with different *LjSRK* haplotypes in weeks 8 and 3, respectively, were highly significant (Paired Student’s t-test, *P <* 0.0001) (Figure 4, A and B).

**Figure 4.**
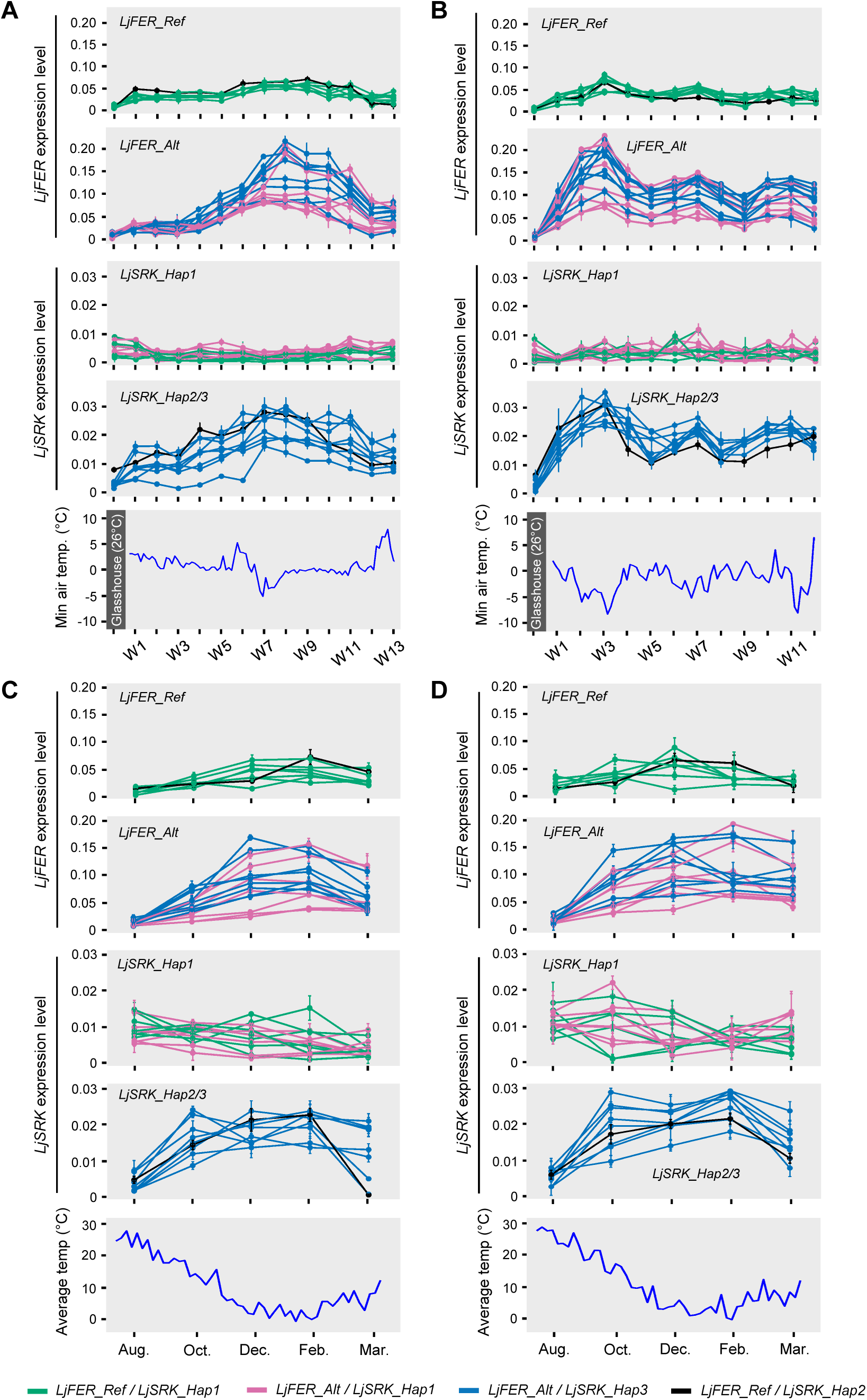
Temperature- and haplotype-dependent *LjFER* and *LjSRK* expression patterns. A-B) Gene expression during winter 2019/20 (A) and winter 2020/21 (B). Leaf samples were collected one week after initiation of outdoor exposure. Glasshouse indicates the sample collected prior to outdoor exposure. C-D) Gene expression levels observed in samples from the Tohoku field site for first year plants (C) and second year plants (D) at five sampling times across the growth season. The leaf samples from first and second year plants were collected from the same plant growing for two consecutive years. Data are presented as mean ± SD of three biological replicates with three technical replicates. The graphs are colored by *LjFER/LjSRK* haplotype combinations. Graphs colored by accession identities are found in Supplemental Figures S11-S12.

Zooming in on sets of accessions originating from similar geographical locations that show *LjFER/LjSRK* haplotype differences, we found consistent patterns of haplotypes, expression and freezing tolerance. From the Kaimon area in Kagoshima MG028 (*LjFER_Ref/LjSRK_Hap1*) consistently showed lower peak *LjFER* expression and lower survival rates than MG022 (*LjFER_Alt/LjSRK_Hap1*), while no differences in *LjSRK* expression were observed (Supplemental Figure S8). Similarly, the accessions from the Aso area showed a gradient of freezing tolerance consistent with the haplotype combinations and expression levels. The *LjSRK_Hap2* haplotype found in MG113 showed clear temperature responsiveness (Supplemental Figure S9).

The glasshouse experiments represent a non cold-acclimated response, where plants are transferred directly from a warm glasshouse into cold and dry winter conditions. The logistic regression results suggested that *LjFER* and *LjSRK* also showed haplotype-dependent seasonal variation. We investigated this further by sampling the same 23 accessions grown in the field across the growth season and quantifying *LjFER* and *LjSRK* expression. The field-grown material showed clear haplotype-dependent differences in seasonal expression profiles (Figure 4, C and D). These were especially clear for *LjSRK*, as most accessions displayed some degree of seasonal variation in *LjFER* expression (Figure 4, C and D), similarly to what we observed for the glasshouse experiment (Figure 4, A and B).

## DISCUSSION

In our earlier study, we combined population structure-corrected GWA and population differentiation analyses to identify traits and associated genes under direct selection during colonization. In-field winter survival under stressful conditions showed both the strongest GWA signals and the most pronounced association with population differentiation (Shah et al., 2020). GWA scan resolution is generally high, with linkage disequilibrium (LD) decaying over a few kb in the Japanese Lotus population sample (Shah et al., 2020). However, it varies with local LD patterns, and selecting specific candidate genes for molecular genetic analysis was not straightforward. Here, we compared the GWA results with differentiation of seasonal transcriptional profiles between northern and southern populations to focus a smaller subset of genes. Although the two chosen candidates, *LjFER* and *LjSRK*, did not show the strongest GWA signals (Figure 1A), they were characterized by both genetic and transcriptional differentiation between populations adapted to different climates and a stronger association with in-field winter survival then could be explained by population structure alone. In addition, the *fer-1* and *srk* mutants showed very dramatic loss of non-acclimated freezing tolerance (Figure 3). Together, these characteristics argue strongly for the relevance of these two receptor-like kinases for Lotus freezing tolerance and climate adaptation.

*LjSRK* resides in a genomic region with six copies of very similar G-type lectin S-receptor-like serine/threonine-protein kinases (Supplemental Figure S10), and the transcription data was helpful in identifying *LjSRK* as the prime candidate. This tandem amplification together with the role of *LjSRK* in climate adaptation suggests positive selection for diversification and sub-or neofunctionalization, and is reminiscent of the amplification of *CBF* genes in Arabidopsis, *M. truncatula*, wheat and barley (Medina et al., 1999; Knox et al., 2010; Tayeh et al., 2013; Würschum et al., 2017). *CBF* genes did not appear as candidates in our GWA analysis. This may be explained by selection mainly acting on non-acclimated freezing tolerance for Lotus, with the *CBF* genes playing a more prominent role in cold acclimation. There is evidence that cold acclimation and *CBF* copy numbers are genetically linked to vernalization in cereal winter annuals (Knox et al., 2010), whereas Lotus does not require vernalization. *CBF* genes could consequently be integrated differently in Lotus cold responses. It is also worth noting that *CBF* genes were not identified as candidates in an Arabidopsis GWA study (Horton et al., 2016), despite their adaptive importance in this annual species (Alonso-Blanco et al., 2005).

In Lotus, the large number of *LjSRK* copies has afforded some flexibility at the protein level, since the haplotypes show amino acid substitutions in addition to their different expression characteristics (Figure 1D). It is striking that although the single instance of *LjSRK_Hap2* shows similar cold responsiveness to *LjSRK_Hap3*, it represents the only example of a cold-responsive *LjSRK* haplotype found together with a *LjFER_Ref* haplotype, whereas all *LjSRK_Hap3* haplotypes are found in *LjFER_Alt/LjSRK_Hap3* combinations. This could either be due to independent selection at the two loci greatly favoring the *LjFER_Alt/LjSRK_Hap3* combination in northern regions or be the result of *LjFER_Ref/LjSRK_Hap3* genetic incompatibility.

In contrast to *LjSRK, LjFER* is a single-copy gene in Lotus. It shows no variation at the protein level, despite presence of coding region SNPs, and the population differentiation is exclusively seen at the regulatory level (Figure 1C). This is consistent with the role of *FER* as a universal and highly conserved growth regulator involved in integration of plant hormone and peptide signaling related to both biotic and abiotic stresses in addition to its essential role in fertility through facilitation of communication between male and female gametophytes (Huck et al., 2003; Liao et al., 2017). Interestingly, *FER* was not identified among 6061 transcripts identified as cold responsive in a large study using ten diverse Arabidopsis ecotypes (Barah et al., 2013), but Arabidopsis *Atfer-4* mutants showed loss of non acclimated freezing tolerance (Chen et al., 2016). This is similar to our observations for Gifu, which does not carry a cold-responsive *FER* haplotype (Figure 2A), but showed a very clear loss of non acclimated freezing on *FER* loss of function (Figure 3, B and C). Lotus recruitment of *FER* into the set of cold-regulated genes, represents an example of recruitment of genes from existing stress-response pathways to promote freezing tolerance, which is hypothesized to be a widespread phenomenon (Preston and Sandve, 2013). In both Lotus and Arabidopsis, *FER* is required for basal non acclimated freezing tolerance. In contrast to annual Arabidopsis, however, *FER* is cold responsive to some degree in all tested perennial Lotus accessions (Figure 3, B-D), but the *LjFER_Alt* haplotype affords a much greater amplitude in the cold and seasonal response, especially to northern accessions that also carry the *LjSRK_Hap3* haplotype (Figure 3, B-D). The *LjSRK_Hap3* haplotype is not required for a strong *LjFER* cold/seasonal response, though, as illustrated by MG110, which carries a *LjFER_Alt/LjSRK_Hap1* haplotype (Supplemental Figures S11-S12).

In summary, we have identified two receptor-like kinases, *LjFER* and *LjSRK*, that mediate non acclimated freezing tolerance and climate adaptation in the perennial herb Lotus. They may represent adaptations specific to perennial herbs, although *SRK* is highly conserved also in annual legumes (Supplemental Figure S13). Because of its central role in growth regulation and extensive conservation, the recruitment of *FER* as a cold-responsive gene in Lotus potentially has broad implications for selecting or engineering frost tolerance in crops.

## MATERIALS AND METHODS

### Read mapping, variant calling and GWA analysis

The previously generated reads (Shah et al., 2020) were mapped to the *L. japonicus* Gifu genome v.1.2 (Kamal et al., 2020), using the *mem* function from the Burrows-Wheeler Aligner (BWA) with default parameters (Li and Durbin, 2009). A mapping quality filter of 30 was applied to the BAM files using SAMtools (Li et al., 2009), which removed reads that mapped to multiple locations. Variant calling was performed on the filtered, sorted and indexed BAM files using *mpileup* and *call* from BCFtools (Danecek et al., 2021). The resulting VCF file included 9,008,653 putative SNPs.

Lotus accessions are highly inbred. The number of heterozygous calls in the VCF file, however, was higher than expected. We inspected the genotype likelihoods and observed numerous inconsistencies on the heterozygous calls. In the variant calling process, especially when the coverage was low, a locus was called as heterozygous even though the genotype likelihood was not 0 for a heterozygous call. For such calls, genotype likelihoods of reference or alternative homozygous were 0 instead. Therefore, we used a custom Python script to correct for these false heterozygous calls. After this step, the number of heterozygous calls was reduced to a reasonable level. Then, a filter for inbreeding coefficient (IBC) was applied (Shah et al., 2020). When the IBC is higher than 0, it indicates that the number of the heterozygotes are lower than the expected according to Hardy-Weinberg principle. This suits our data well, thus we only took the loci where IBC > 0. The other criteria that we used in our filtering step were as follows: (1) mapping quality (MQ) > 30, (2) total depth (DP) > 60, (3) quality (QUAL) > 60, (4) must be biallelic, (5) missing genotypes < 50%, (6) Gifu and MG-20 must be homozygous, and (7) minor allele frequency (MAF) > 0.05.

GWA analysis was carried out using the approach previously described (Shah et al., 2020). Prior to GWA analysis, missing genotype data points were imputed using Beagle (Browning et al., 2018) and the VCF file was converted to a simple genotype-only format (012 format) with VCFtools (Danecek et al., 2011). Then GWA analysis was performed using a Python implementation of EMMA (https://app.assembla.com/spaces/atgwas/git/source) (Kang et al., 2008; Atwell et al., 2010) with a minor allele count cutoff of 10. Finally, the Manhattan plot was made using the *qqman* package in R (Turner, 2018).

### Plant materials and field phenotyping

To investigate the plant winter-survival phenotype under field conditions, we grew the Lotus accessions at a field site located near the Graduate School of Life Sciences, Tohoku University, Kashimadai, Miyagi Prefecture, Japan (38.46°N, 141.09°E). All seeds were scarified and sterilized with 2% of NaClO then were transferred to 0.8% agar plate and incubate in a controlled growth chamber for three days of dark and two days for normal condition (70% HR with 16 h : 8 h, light : dark at 25°C). Germinated seeds were transferred into plant cell trays containing 1:1 mixture of soil and vermiculite, and were grown in a glasshouse for one month before transplanting to the field. Two individuals from the same accessions were transplanted, spaced least 50 cm apart directly into the field soil. These sets of two individuals per accession were planted with 60 cm intervals. Each accession consisted of eight sets of replicates, a total 16 plants grown per accession. The transplanting was carried out in early May 2018 and plants were then grown for two consecutive years to the end of March 2020. Leaf samples from three plants of each accession were harvested at five time points (August, October, December, February and March) for gene expression analysis in two consecutive years. The field air temperature data were obtained from the Automated Meteorological Data Acquisition System (AMeDAS) in Kashimadai provided by Japan Meteorological Agency (https://www.jma.go.jp/jp/amedas), as AMeDAS Kashimadai was located at the Tohoku field. The winter survival rate was calculated as the ratio between the total number of surviving plants (observed during early spring) to the total number of plants growing before onset of winter (observed during early fall).

### Transcriptional profiling and logistic regression analysis

To facilitate disentangling geographic origin and population structure, we carried out transcriptome profiling of selected accessions (Figure 1B). Total RNA from leaf samples collected in August, October, December and February 2018/2019 was extracted using the RNeasy Plant Mini kit (Qiagen) according to the manufacturer’s instructions. The residual genomic DNA contaminants were then removed with DNase I treatment (Qiagen). The mRNA-seq libraries were constructed by using SureSelect Strand Specific RNA Library Prep kit (Agilent Tech.) following the manufacturer’s recommendations. The libraries were then sequenced on a NextSeq 500 (Illumina Inc.) with 75 base single-end reads. The reads were mapped to the *L. japonicus* Gifu genome (v.1.2) (Kamal et al., 2020) using the Bowtie2 (v.2.3.5) aligner (Langmead and Salzberg, 2012) and the number of reads mapped per gene was quantified using SAMtools (v.1.9) (Li et al., 2009). The resulting expression profiles were used in logistic regression analysis (Supplemental Table S3). For this, the accessions were split into two groups based on latitude and pop3 membership, respectively. The accessions were assigned to one group if their latitude or pop3 membership was larger than the mean and to the other if their value was lower than the mean. Logistic regression analysis, as implemented in the Python library *sklearn*, was then used for accession classification into the two groups based on the expression profile of a given gene, and the fraction of correctly classified accessions was recorded for each gene for both latitude and pop3 membership.

### Outdoor winter challenge experiment

Scarified and sterilized treated seeds were germinated in 0.8% agar followed by transferring into a 1:1 mixture of soil and vermiculite. The plants then grew for four weeks in the glasshouse at ∼26°. Thirteen to sixteen individuals of each accession were used. Afterwards, the plants were moved to the open field for four-weeks during the winter season, exposing them to natural winter conditions. During snowy days, we did not remove the snow that covered the plant surface. Similarly, the frozen water surrounding the plants was not removed during the freezing day. The daily minimum air temperatures were recorded using a digital thermometer during the exposure periods. After four weeks of exposure, the plants were returned to the glasshouse for recovery at ∼26° for two weeks. Leaf samples from three plants were collected on the day before treatment (BT) inside the glasshouse and after one of winter exposure (day-7). This set of experiments was repeated weekly from early December to late March in winter 2019/20 and winter 2020/21. The outdoor temperatures for the two experimental years are shown in Figure 2 and Figure 4.

### *LORE1* functional analysis

To evaluate the function of the candidate genes, the Lotus retrotransposon (*LORE1*) insertion tag lines were identified using *Lotus* Base (https://lotus.au.dk/lore1/search) (Mun et al., 2016), and used a single insertion line in *LjFER* with ID 30049953 as *fer-1* and 3 insertion lines in *LjSRK* with mutant IDs 30124875, 30057969 and 30007797 as *srk-1, srk-2* and *srk-3*, respectively. All *LORE1* lines are in the Gifu genetic background (Małolepszy et al., 2016). The *LORE1* and Gifu seeds were treated and germinated as described for the Japanese accessions above. *LORE1* genotyping was performed by “touchdown PCR” using the PCR primers provided by *Lotus* Base (Mun et al., 2016) to confirm the insertion integrity as well as the homozygosity in the *LORE1* lines using Gifu as the wild type control. Primer sequences are listed in Supplemental Table S7 (Urbański et al., 2012; Mun et al., 2016). The gene expression of the target genes was then analyzed in all these plants including the non-insertion and heterozygous individual before transferring to either field or glasshouse, using the same procedures as described for the Japanese wild accessions. For the outdoor winter exposure, we harvested leaf samples from six *LORE1* mutant plants as well as Gifu on the day before treatment (BT) inside the glasshouse, and at the end of week-1 (W1) and week-3 (W3) of winter exposure for gene expression analysis. Recovery took place as described for the Japanese wild accessions. This set of experiments was carried out in two experimental years, winter 2019/20 and winter 2020/21. The winter 2020/21 experiment was performed in three experimental replicates. The periods as well as the observed minimum air temperature of outdoor winter exposure are shown in Supplemental Figure S2, A and B.

### RNA extraction and gene expression analysis by qRT-PCR analysis

*LjFER* and *LjSRK* gene expression was quantified using real-time quantitative PCR. Total RNA from leaf samples collected during field experiments, outdoor winter challenges, and *LORE1* analysis was extracted using the RNeasy Plant Mini kit (Qiagen) and was treated with DNase I (Qiagen). cDNA was synthesized from total RNA using PrimeScript™ reverse transcriptase reagent Kit (Perfect Real Time) (Takara) according to the manufacturer’s instructions. Three and six biological replicates of Japanese accessions and *LORE1* mutants were used for the analysis, respectively. Real-Time qPCR was performed using KAPA SYBR® FAST qPCR Master Mix (KAPABiosystems) on the CFX Connect Real-time PCR system (Bio-rad) according to the manufacturer’s instructions. *LjFER-* and *LjSRK-*specific primers were designed using Primer3Plus (sequences listed in Supplemental Table S7) (Untergasser et al., 2012). The gene expression level was determined by comparison to the reference gene *LjUbiq* using the formula 2^-ΔCT^ where ΔCT is the crossing point difference between target and reference gene. Three technical replicates for each accession and target gene were conducted.

## Statistical analysis

The statistical analysis was performed using R software with its own function (v.3.6.1). The haplotype comparison for winter survival rate in 2014 (Figure 1, E and F) was analyzed using one-way ANOVA followed by posthoc Tukey’s test in R package *agricolae*. The comparisons for outdoor winter challenge recovery boxplots (Figure 2D) and gene expression by Real-Time qPCR were analyzed using Paired Student’s t-test. In the figures, the statistical significance represents by the following notations: ns = no significant (*P* > 0.05), * = *P* < 0.05, ** = *P* < 0.01, *** = *P* < 0.001, **** = *P* < 0.0001.

## Code availability

The code used in this study is available on GitHub in the repository https://github.com/yusdarmust/Lotus_overwintering.

## Accession numbers

The RNA-seq data used in this study have been deposited and available in DDBJ Read Sequence Archive repository under accession numbers DRR342836-DRR342883.

## SUPPLEMENTAL INFORMATION

**Supplemental Figure S1**. *LORE1* mutant in Tohoku field.

**Supplemental Figure S2**. Observed air temperature of *LORE1* mutant outdoor exposure in two experimental years.

**Supplemental Figure S3**. *LjFER* and *LjSRK* expression in *LORE1* mutant during outdoor exposure of winter 2019/20.

**Supplemental Figure S4**. *LjFER* expression in Gifu wild type and *fer-1* heterozygous lines before outdoor exposure of winter 2020/21.

**Supplemental Figure S5**. Recovery rates of *fer* and *srk LORE1* mutants during winter exposure 2020/21.

**Supplemental Figure S6**. *LjFER* and *LjSRK* expressions in *LORE1* mutants during outdoor winter exposure 2020/21.

**Supplemental Figure S7**. The phenotypes of *fer* and *srk LORE1* mutants as well as Gifu WT during outdoor winter exposure 2020/21.

**Supplemental Figure S8**. Exposure survival and expression of the accessions from Kaimon area, Kagoshima during two years of outdoor winter exposure.

**Supplemental Figure S9**. Exposure survival and expression of the accessions from the Aso area during two years of outdoor winter exposure.

**Supplemental Figure S10**. Genomic region where *LjSRK* resides with other very similar G-type lectin S-receptor-like serine/threonine-protein kinases.

**Supplemental Figure S11**. Temperature- and haplotype-dependent *LjFER* and *LjSRK* expression patterns during outdoor winter exposure with accession-highlight.

**Supplemental Figure S12**. Temperature- and haplotype-dependent *LjFER* and *LjSRK* expression patterns in Tohoku field in the 1st and 2nd year plant with accession-highlight.

**Supplemental Figure S13**. Multiple alignment of *LjSRK* with its orthologues in annual legumes.

**Supplemental Table S1**. Genotype calls based on the reference assembly

**Supplemental Table S2**. GWAS results for overwintering 2014 phenotype based on Gifu reference assembly.

**Supplemental Table S3**. RNA-seq expression data based on RPKM value. **Supplemental Table S4**. Logistic regression results.

**Supplemental Table S5**. Fifty-nine SNPs corresponded with 12 GWAS candidate genes which intersect with the candidate genes from the logistic regression analysis.

**Supplemental Table S6**. Observed minimum air temperature (°C) and accession recovery rates of outdoor winter exposure.

**Supplemental Table S7**. PCR primers used for *LORE1* genotyping and Real-time PCR.

## ACKNOWLEDGMENTS

We thank C. Mitsuoka and M. Chiba for excellent technical assistance. Accessions of *L. japonicus* were provided by the National BioResource Project ‘*Lotus*/*Glycine*’.

## AUTHOR CONTRIBUTIONS

S.S. and S.U.A. conceived, coordinated and supervised the project; S.S., S.U.A. and Y.M. designed the experiments; Y.M. performed *LORE1*, outdoor winter exposure, and gene expression experiments; T.Y.A. performed variant calling and GWA analysis; M.G. and C.W. performed the transcriptional profiling; Y.M., T.Y.A. and M.B. analyzed the data; A.M.M., Y.I. and Y.K. performed all other experiments; S.S. and S.U.A. provided the resources; S.U.A., Y.M. and S.S. wrote the paper with input from all authors.

## DATA AVAILABILITY

All data are either in the main paper or the Supporting Information. Source data are provided with this paper. Requests should be directed to the corresponding authors. The Supplemental Table S1 was deposited and publicly available on Mendeley Data repository with DOI: 10.17632/yxhz58mx73.1 titled “GWA OW - Genotype calls”.

